# *ent*-Verticilide B1 inhibits type 2 ryanodine receptor channels and is antiarrhythmic in Casq2-/- mice

**DOI:** 10.1101/2023.07.03.547578

**Authors:** Aaron Gochman, Tri Q. Do, Kyungsoo Kim, Jacob A. Schwarz, Madelaine P. Thorpe, Daniel J. Blackwell, Abigail N. Smith, Wendell S. Akers, Razvan L. Cornea, Derek R. Laver, Jeffrey N. Johnston, Bjorn C. Knollmann

**Author notes:** Corresponding author: Bjorn C. Knollmann, MD, PhD Medical Research Building IV, Rm. 1265 2215B Garland Ave Nashville, TN 37232-0575.

## Abstract

Ca^2+^ leak from cardiac ryanodine receptor (RyR2) is an established mechanism of sudden cardiac death (SCD), whereby dysregulated Ca^2+^ handling causes ventricular arrhythmias. We previously discovered the RyR2-selective inhibitor *ent-*(+)-verticilide (*ent*-1), a 24-membered cyclooligomeric depsipeptide that is the enantiomeric form of a natural product (*nat*-(-)-verticilide). Here, we examined its 18-membered ring-size oligomer (*ent*-verticilide B1; “*ent*-B1”) in single RyR2 channel assays, [^3^H]ryanodine binding assays, and in *Casq2*^-/-^ cardiomyocytes and mice, a gene-targeted model of SCD. *ent*-B1 inhibited RyR2 single-channels and [^3^H]ryanodine binding with low micromolar potency, and RyR2-mediated spontaneous Ca^2+^ release in Casq2-/- cardiomyocytes with sub-micromolar potency. *ent*-B1 was a partial RyR2 inhibitor, with maximal inhibitory efficacy of less than 50%. *ent*-B1 was stable in plasma, with a peak plasma concentration of 1460 ng/ml at 10 min and half-life of 45 min after intraperitoneal administration of 3 mg/kg in mice. Both 3 mg/kg and 30 mg/kg *ent*-B1 significantly reduced catecholamine-induced ventricular arrhythmia in Casq2-/- mice. Hence, we have identified a novel chemical entity – *ent*-B1 – that preserves the mechanism of action of a hit compound and shows therapeutic efficacy. These findings strengthen RyR2 as an antiarrhythmic drug target and highlight the potential of investigating the mirror-image isomers of natural products to discover new therapeutics.

**Significance statement:** The cardiac ryanodine receptor (RyR2) is an untapped target in the stagnant field of antiarrhythmic drug development. We have confirmed RyR2 as an antiarrhythmic target in a mouse model of sudden cardiac death and shown the therapeutic efficacy of a second enantiomeric natural product.

## Introduction

Patients with heart rhythm disorders have a major, unmet need for new treatments. Among causes of mortality in the United States, sudden cardiac death (SCD) – a result of ventricular arrhythmias – is responsible for 15% of all deaths annually (Zheng et al., 2001). With the exception of beta-adrenergic receptor inhibitors, none of the agents marketed as antiarrhythmic drugs in the U.S. prevent SCD, with most of them increasing rates of mortality in patients with ischemic heart disease or heart failure (Cardiac Arrhythmia Suppression Trial, 1989; Kober et al., 2008; Waldo et al., 1996). The common mechanism of action shared by FDA-approved antiarrhythmic drugs is the modulation of ion channels or G-protein coupled receptors expressed in the cell membrane (Brunton and Knollmann, 2023). As such, better antiarrhythmic drugs are needed. Here we build on our previous work (Batiste et al., 2019; Kryshtal et al., 2021) to help establish a new class of antiarrhythmic drugs by further validating the intracellular target of cardiac ryanodine receptors (RyR2) for antiarrhythmic drug development.

RyR2s are Ca^2+^ release channels located in the membrane of the sarcoplasmic reticulum (SR). The mechanism of Ca^2+^-induced Ca^2+^ release to facilitate excitation-contraction (EC) coupling in cardiomyocytes is well-studied, with RyR2 serving as the release channel for SR Ca^2+^ stores (Bers, 2002). Pathologic Ca^2+^ release from RyR2 has been reported in both genetic and acquired arrhythmia disorders through gain of function mutations (Watanabe and Knollmann, 2011) or post-translational modifications to RyR2 (Marx et al., 2000; Respress et al., 2012; Terentyev et al., 2008), respectively. In either scenario, an increased open probability of RyR2 causes Ca^2+^ to “leak” from the SR, which disrupts both the temporal and functional integrity of cardiac Ca^2+^ handling. Specifically, increased [Ca^2+^]_cytosolic_ is pumped extracellularly through the electrogenic Na^+^/Ca^2+^exchanger, leading to delayed afterdepolarizations driven by the untimely influx of Na^+^ as Ca^2+^ homeostasis is restored (Knollmann and Roden, 2008).

To address the need for a first-in-class RyR2-selective drug, we discovered a bioactive enantiomer of the fungal natural product verticilide (Shiomi et al., 2010). This hit compound, called *ent-*(+)-verticilide (*ent-*1), is a cyclooligomeric depsipeptide (COD) that selectively inhibits RyR2 *in vitro* (Batiste et al., 2019). Unlike *nat*-(-)-verticilide, *ent*-1 therapeutically inhibited RyR2 Ca^2+^ leak in mice with catecholaminergic polymorphic ventricular tachycardia (CPVT), a genetic disease that can present as exercise- or stress-induced SCD in childhood (Leenhardt et al., 1995). While *ent*-verticilide has become a useful tool compound in exploring the potential of RyR2-targeted therapy in animal models of arrhythmia (Kim et al., 2023), a follow-up structure-activity study revealed that the oligomeric structure of *ent*-1 can be modified to an alternate ring-size analog that maintains an affinity for RyR2 (Smith et al., 2021). The 18-membered ring analog *ent*-verticilide B1 (*ent*-B1) (MW = 639.8 g/mol; *ent*-1 MW = 853.1 g/mol) retained RyR2 inhibitor activity in a Ca^2+^ spark assay using permeabilized cardiomyocytes, an accepted measure of isolated RyR2 Ca^2+^ release (Smith et al., 2021). *ent*-B1 is also the mirror image of a natural product, verticilide B1, that is not active against RyR2 or insect RyR (Ohshiro et al., 2012). Here, we studied the *in vitro* pharmacology of *ent*-B1 and tested the hypothesis that *ent*-B1 has antiarrhythmic efficacy *in vivo*.

## Material and Methods

### Drugs, chemicals, and reagents

All chemicals and reagents were purchased from Sigma-Aldrich unless otherwise stated.

### Single Channel Recording

SR vesicles containing RyR2 were isolated from sheep hearts and incorporated in artificial bilayer membranes as previously described (Laver et al., 1995). Lipid bilayers were formed across an aperture with diameter 150-250 mm of a delrin cup using a lipid mixture of phosphatidylethanolamine and phosphatidylcholine (8:2 wt/wt, Avanti Polar Lipids, Alabaster, AL) in n-decane (50 mg/ml, ICN Biomedicals, Irvine, CA). During the SR vesicle fusion period, the cis (cytoplasmic) chamber contained 250 mM Cs^+^ (230 mM CsCH_3_O_3_S, 20 mM CsCl) + 1.0 mM CaCl_2_ and the trans (luminal) chamber contained 50 mM Cs^+^ (30 mM CsCH_3_O_3_S, 20 mM CsCl) + 1 mM CaCl_2_. When ion channels were detected in the bilayer, the trans Cs^+^ was raised to 250 mM by aliquot addition of 4 M CsCH_3_O_3_S. During experiments, the cis solution was exchanged by a perfusion system (O’Neill et al., 2003) to one containing 250 mM Cs^+^ plus 2 mM ATP and free Ca^2+^ of 100 nM followed by exchange with the same plus *ent*-B1. Thus, the perfusion system allowed repeated application and washout of *ent*-B1 within ∼3 s.

All solutions were pH buffered using 10 mM TES (N-tris[hydroxymethyl] methyl-2-aminoethanesulfonic acid; ICN Biomedicals) and titrated to pH 7.4 using CsOH (ICN Biomedicals). Free Ca^2+^ of 100 nM was generated from 1 mM CaCl_2_ and 4.5 mM BAPTA (1,2-bis(o-aminophenoxy)ethane-N, N, N’, N’-tetraacetic acid; obtained from Invitrogen) and this was validated using a Ca^2+^ electrode (Radiometer, Brea, CA). ATP was in the form of the di-sodium salt and obtained from Enzo Life Sciences (Farmingdale, NY) and Cs^+^ salts were obtained from Sigma-Aldrich (St Louis, MO). CaCl_2_ was obtained from BDH Chemicals (VWR, Radnor, PA). Cytoplasmic recording solutions were buffered to a redox potential of -232 mV with glutathione disulfide (GSSG; 0.2 mM) and glutathione (GSH; 4 mM; MP Biomedicals), and luminal solutions were buffered to a redox potential of -180 mV with GSSG (3 mM) and GSH (2 mM), both obtained from MP Biomedicals. *ent*-B1 was prepared as a stock solution in DMSO.

### Single Channel Recording Analysis and Data Acquisition

Experiments were carried out at room temperature (23 ± 2 °C). Electric potentials are expressed using standard physiological convention (*i.e.* cytoplasm relative to SR lumen at virtual ground). Control of the bilayer potential and recording of unitary currents was done using an Axopatch 200B amplifier (Axon Instruments/Molecular Devices, Sunnyvale, CA). Channel currents were digitized at 50 kHz and low pass filtered at 5 kHz. Before analysis the current signal was redigitized at 5 kHz and low pass-filtered at 1 kHz. Individual readings of open probability were derived from 30-60 s of RyR2 recording. Single channel open probability was measured using a threshold discriminator at 50% of channel amplitude. *[^3^H]ryanodine ligand binding assay* – Isolated cardiac SR vesicles were incubated with, 200 nM CaM binding peptide, 0.1 µM CaCl2, 20 mM PIPES, 150 mM KCl, 5mM GSH, 0.1 mg/mL BSA, 1 µg/mL aprotininin, 1µg/mL leupeptin, and 1 µM DTT for 30 min at 37 °C. Samples were centrifuged at 110,000 x g for 25 min at 4 °C and resuspended to at final concentration of 15 mg/mL in 20 mM PIPES, 150 mM KCl, 5mM GSH, 0.1 mg/mL BSA, 1 µg/mL aprotinin, 1µg/mL leupeptin, and 1 µM DTT. In 96-well plates, cardiac SR membranes (CSR, 3 mg/mL) were pre-incubated with 1% v/v *ent*-B1 or *ent*-1 (to yield the indicated drug concentrations) for 30 min, at 22 °C, in a solution containing 150 mM KCl, 5 mM GSH, 1 µg/mL Aprotinin/Leupeptin, 1 mM EGTA, and 23 µM or 1.62 mM CaCl_2_ (100 nM or 30 µM free Ca^2+^, respectively as determined by MaxChelator), 0.1 mg/mL BSA, and 20 mM K-PIPES (pH 7.0). Non-specific [^3^H]ryanodine binding to SR was assessed by addition of 15 µM non-radioactive ryanodine. Maximal [^3^H]ryanodine binding was assessed by addition of 5 mM adenylyl-imidodiphosphate (AMP-PNP), supplemented with 20 mM caffeine. These control samples were each loaded over four wells per plate. Binding of [^3^H]ryanodine (7.5 nM) was determined following a 3 h incubation at 37 °C and filtration through grade GF/B glass microfiber filters (Brandel Inc., Gaithersburg, MD, US) using a M96T-Brandel Harvester. Filters were immersed in 4 mL of Ecolite scintillation cocktail and incubated 24 hours prior to [^3^H] counting in a Perkin-Elmer Tri-Carb 4810.

### Intracellular Ca^2+^ measurements in intact cardiomyocytes

Cardiomyocytes were pre-incubated for 2 hours with DMSO, *ent*-1, or *ent*-B1. Myocytes were then loaded with Fura-2 acetoxymethyl ester (Fura-2 AM; Invitrogen) as described previously (Batiste et al., 2019). Briefly, isolated single ventricular myocytes were incubated with 2 µM Fura-2 AM for 7 minutes to load the indicator in the cytosol. Myocytes were then washed twice for 10 minutes with normal Tyrode (NT) solution containing 250 µM probenecid (all solutions contained DMSO, *ent*-1, or *ent*-B1). The composition of NT used for Fura-2 loading and washing was (in mM): 134 NaCl, 5.4 KCl, 1.2 CaCl2, 1 MgCl2, 10 glucose, and 10 HEPES, pH adjusted to 7.4 with NaOH. After Fura-2 loading, all experiments were conducted in NT solution containing 1 µM isoproterenol and 2 mM CaCl2. Fura2-loaded myocytes were electrically paced at 3 Hz field stimulation and Ca transients were recorded for 20 seconds followed by no electrical stimulation for 40 seconds to record spontaneous Ca release events. After that, myocytes were perfused with 10 mM caffeine in NT solution for 5 seconds to estimate total SR Ca content. Fura-2 was measured using a dual-beam excitation fluorescence photometry setup (IonOptix Corp.) and analyzed using commercially available data analysis software (IonWizard, IonOptix, Milton, MA). All experiments were conducted at room temperature.

### In vivo pharmacokinetic study

The pharmacokinetic study was carried out by a contract research organization, Pharmaron, Inc. Mice were housed with free access to food and water. All protocols and procedures were compliant with Animal care and Use Application approved by the Institution Animal Care and Use Committee of Pharmaron, Inc., following the guidance of the Association for Assessment and Accreditation of Laboratory Animal Care (AAALAC). Based on our previous study with *ent*-1 (Blackwell et al., 2023), an *en*t-B1 dose of 3 mg/kg (drug/body weight) was used. Three CD1 mice (all males) 6-8 weeks old were selected for the study. Mice were injected intraperitoneally with *ent*-B1 dissolved in solution containing 10% Tween 20, 10% DMSO, 40% water, and 40% PEG-400 (v/v, 5 mL/kg), with a final concentration of 0.6 mg/mL. 30 µL of blood was collected from each animal at 10, 20, 30, 60, 180, and 480 minutes following drug administration and centrifuged at 5000 x g, 4 °C for 5 minutes to obtain plasma. The samples were stored at -75 ± 15 °C until analysis. Clinical observation showed no abnormality during the entire experiment.

### LC-MS/MS analysis of ent-B1

LC-MS/MS with electronspray ionization in the positive ion mode setting was used to detect *ent*-B1 followed by multiple reaction monitoring of precursor and product ions as follows: *ent*-B1 (mass-to-charge ratio [m/z] 640.18 to 214.00). Mouse plasma was quantified using nine standards (0.5 – 1000 ng/ml) and four quality control levels (1, 2, 50, 800 ng/ml) prepared independently of those used for the standard curve. 20 µL of standards, quality control samples, and unknown samples (10 µL plasma and 10 µL blank solution) were added to 200 µL acetonitrile containing internal standard (dexamethasone; [m/z] 393.40 to 373.30) for protein precipitation. The samples were vortexed for 30 seconds, centrifuged for 15 minutes at 4000 rpm and 4 ׄ°C, and the supernatant was diluted five times with water. Samples were then loaded into SIL-30AC autosampler and 10 µL was injected into a Shimadzu LC-30AD Series HPLC coupled to an AB Sciex Triple Quad 5500 mass spectrometer. Analytes were separated on a Raptor Biphenyl column (50 x 2.1 mm, 2.7 µm) using a 95:5 (v/v) mobile phase mixture of 0.1% formic acid in water (mobile phase A) and 0.1% formic acid in acetonitrile (mobile phase B) at the flow rate of 0.6 ml/min. The gradient for mobile phase B was increased to 95% over 1.4 minutes with a total run time of 2.5 minutes for each sample. All quality controls and standards met the following acceptance criteria 1) standard curve of at least five standards are within 15% of their nominal concentrations and at least 50% at each quality control level (low, medium, and high) were within 15% of their nominal concentrations.

### Pharmacokinetic analysis

Plasma concentrations of *ent*-B1 at each time point were imported to Phoenix WinNonlin® 8.0 software (Certara USA, Inc., Princeton, NJ). Plasma concentration time profiles for individual animals were analyzed by noncompartmental analysis using model 200 (Plasma; Single Extravascular Dose; Linear Log Trapezoidal Method) to approximate the elimination rate constant (ke), half-life (T_1/2_), maximum observed plasma concentration (C_max_), time to maximum observed plasma concentration (T_max_), the area under the plasma concentration-time curve from zero to infinity (AUC_inf_). Dose was normalized to 3 mg/kg for each animal and used to derive estimates of extravascular clearance (Cl/F) and extravascular volume of distribution (Vz/F) by noncompartmental analysis.

### ECG recording

Nine male and thirteen female *Casq2^-/-^* mice 17-20 weeks old were randomly assigned to a three-by-three crossover design such that every treatment sequence was sampled. Mice were pretreated with intraperitoneal injection of vehicle (DMSO), 3 mg/kg *ent*-B1, or 30 mg/kg of *ent*-B1 fifteen minutes prior to baseline ECG recording. Mice were then injected with 3 mg/kg isoproterenol intraperitoneally. Recordings were continued for 5 minutes or for 1 minute after cessation of ventricular ectopy. A washout period of one week was used between treatments.

### ECG analysis

LabChart (AD Instruments, Inc) was used to analyze ECG recordings by a reviewer blinded to treatment dose. ECG records were examined to quantify premature ventricular contractions (PVCs), duration of ventricular tachycardia, heart rate, and QRS width. Arrhythmias were scored on a five point ordinal scale based on the number of PVCs with the following criteria: 1) zero point for no PVCs; 2) one point for isolated PVCs; 3) two points for bigeminy (alternating sinus beats and PVCs); 4) three points for couplets (two consecutive PVCs); and 5) four points for three or more consecutive PVCs (ventricular tachycardia).

### Statistical Analysis

Statistics were carried out in GraphPad Prism (v9.5.0) or Matlab as indicated in the figure legends. All tests were two-sided and a p-value cutoff of <0.05 was used to denote significance. Concentration-response curves for [^3^H]ryanodine ligand binding assay and Ca^2+^ spark assays were generated using non-linear regression with the equation: Y = Bottom + (Top-Bottom)/(1+(IC_50_/X)^HillSlope) using least square regression, no weighting, and no constrains on the parameters. The distributions of relative RyR2 open probability at each [ent-B1] were normalized by taking the log of each sample. The fitted concentration-response curve was constrained for HillSlope = 1. For the 3 x 3 crossover study, a pre-test was conducted using a mixed model with fixed effects of sequence and period; mice were treated as random effects. When warranted, a post-hoc test was conducted and used, as reported in the pertaining figure legends, to test the null hypothesis that the 3 mg/kg or 30 mg/kg doses do not deviate from treatment with vehicle.

## Results

### ent-B1 inhibits RyR2 single channels and [3H]ryanodine binding in SR vesicles

To determine the effect of *ent*-B1 on RyR2 channel activity, we studied isolated SR vesicles derived from ovine hearts. RyR2 channels were incorporated into artificial lipid bilayers and exposed to *ent*-B1 via perfusion (**Fig. 1**). We tested the effect of three concentrations of *ent*-B1 (10, 30, and 50 µM) on RyR2 channel open probability. Cytosolic Ca^2+^ concentration was kept at 100 nM to match physiological concentrations during diastole. *ent*-B1 inhibited RyR2 channels in a concentration-dependent manner, with an estimated IC_50_ of 25 ± 23 µM and incomplete maximal channel inhibition (**Fig. 1C**). There was a corresponding shift in the current amplitude histogram toward *I*=0 in the presence of *ent*-B1, with no detectable substates. (**Fig. 1D**). To establish a more detailed concentration-response relationship, we performed a [^3^H]ryanodine binding assay using sarcoplasmic reticulum (SR) samples from porcine cardiac muscle. In this assay of RyR2 channel activity, the presence of an inhibitor reduces the amount of [^3^H]ryanodine bound to the samples. When compared to vehicle (DMSO), *ent*-B1 reduced [^3^H]ryanodine binding with a low micromolar potency (IC_50_ = 1.9 μM) and incomplete maximal inhibition of approximately 30% (**Fig. 2**). Taken together, these results demonstrate that *ent*-B1 directly inhibits RyR2 channels.

**Figure 1:**
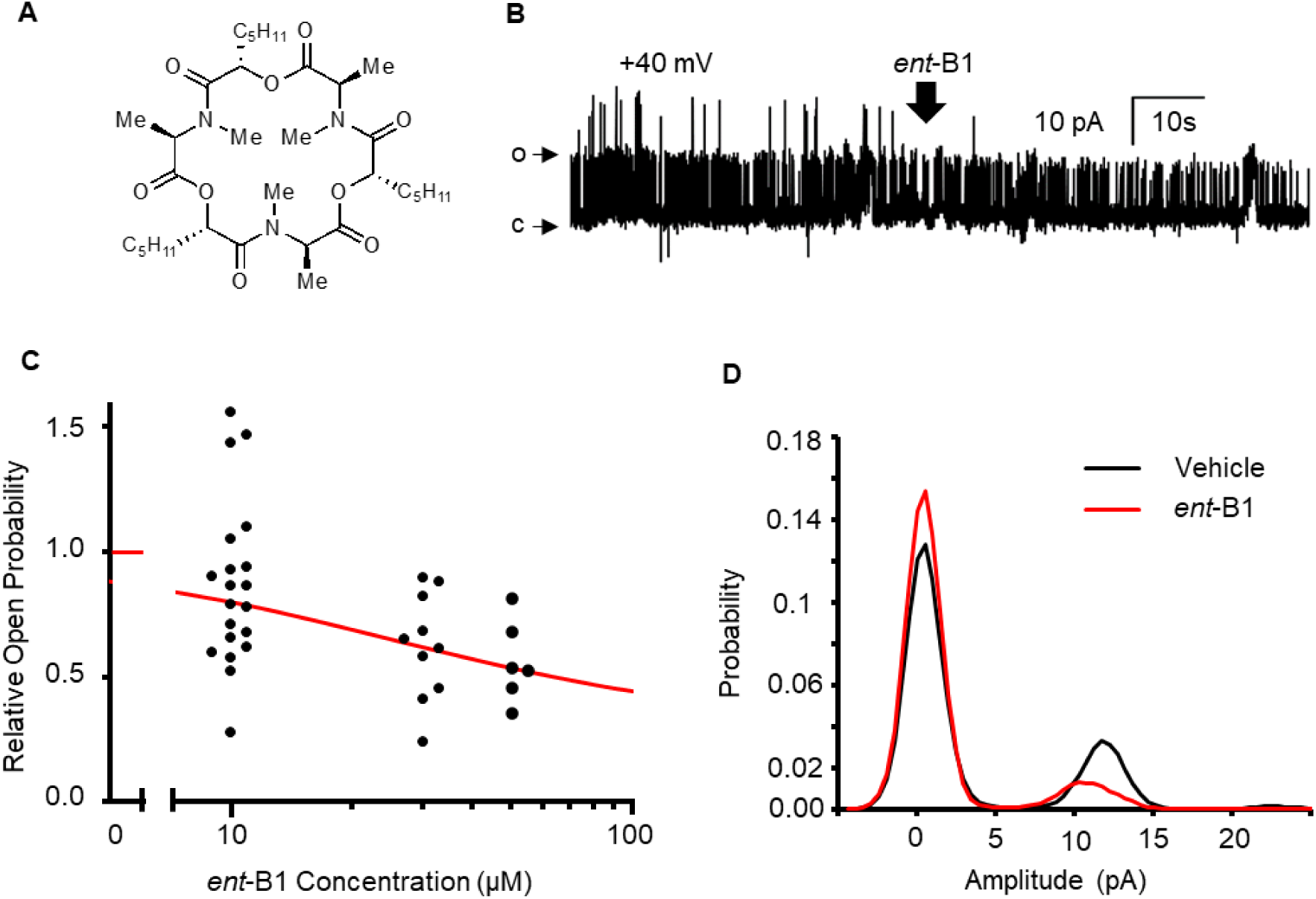
RyR2 single channel recording in artificial lipid bilayers. **A)** Chemical structure of *ent-*B1. **B)** Representative trace of RyR2 single channel recording before and after application of *ent-*B1 at +40 mV, with 2 mM ATP and 100 nM Ca (cis) and 1 mM Ca (trans). Channel openings are in the upward direction (c, closed; o, open). Arrow indicates addition of *ent-*B1. **C)** Open probability of RyR2 in the presence of *ent-*B1 relative to vehicle (DMSO). Fitting values using non-linear regression to a Hill-function (Matlab, red line) yielded an IC_50_ of 25 ± 23 µM and a maximum inhibitory effect (I_max_) of 70 ± 30%. A two-sided sign test (Matlab) determined inhibition at each concentration with *P* = 0.12, 0.002 and 0.031, respectively, compared to vehicle. **D)** Probability distributions of current amplitude of 30-second segments of recording from (B) before and after addition of 30 µM *ent-*B1.

**Figure 2:**
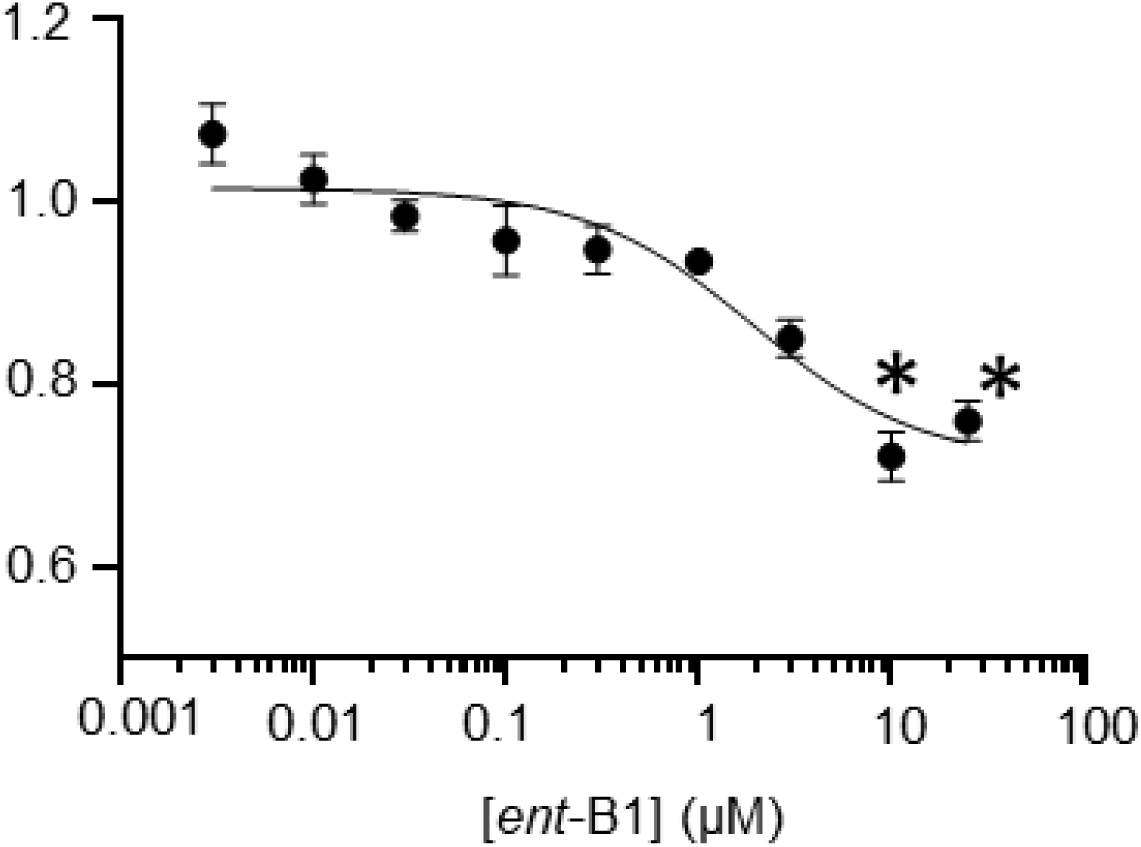
Effect of *ent-*B1 on RyR2 activity in porcine SR vesicles. RyR2 activity was measured by [^3^H]ryanodine binding. The concentration-response curves show results for *ent-*B1 relative to control (DMSO). *ent*-B1 IC_50_ = 1.93 μM (95% CI 0.740-4.62 μM), *I*_max_ = 0.71. Data are shown as mean ± SE (n = 6). \**P*<0.05 vs DMSO by unpaired, two-tailed *t* test

### ent-B1 reduces spontaneous Ca2+ release in intact mouse CPVT cardiomyocytes

Direct pharmacological block of RyR2 necessitates a compound permeate the cell membrane (sarcolemma). Single channel and [^3^H]ryanodine assays do not require cell membrane permeability or transport. Hence, we next examined the activity of *ent*-B1 in cardiomyocytes isolated from *Casq2^-/-^* mice to determine whether *ent*-B1 blocks RyR2 in intact cells. To quantify drug efficacy, we measured the rate of spontaneous RyR2-mediated spontaneous SR Ca^2+^ release (SCR) events following a 3 Hz pacing protocol (**Fig. 3A**). Compared to vehicle control (DMSO), *ent*-B1 reduced the rate of SCRs in a concentration-dependent manner (IC_50_ = 0.23 ± 0.1 µM) with ∼50% efficacy at the highest concentration tested (Fig. 3B). This indicates that *ent*-B1 significantly inhibits pathologic Ca^2+^ release, with higher potency than the single channel and [^3^H]ryanodine binding assays suggested. Only the highest concentration of *ent*-B1 tested (3 µM) reduced diastolic Ca^2+^ levels, the amplitude of the systolic Ca^2+^ transient, and the amplitude of the caffeine-induced Ca^2+^ transient (**Fig. S1**), which is consistent with a mechanism of action of a partial RyR2 inhibitor. *ent*-B1 had no effect on other measures of SR Ca^2+^ release such as time-to-peak of Ca^2+^ transients or Ca^2+^ decay kinetics (**Fig. S1C, D**), the latter being an indicator of SERCA2 Ca^2+^ SR uptake rate. We also saw no effect of *ent*-B1 on the decay rate of the caffeine-induced SR Ca^2+^ transient (**Fig. S1F**), which is a standard measure of the Na^+^-Ca^2+^ exchanger activity.

**Figure 3:**
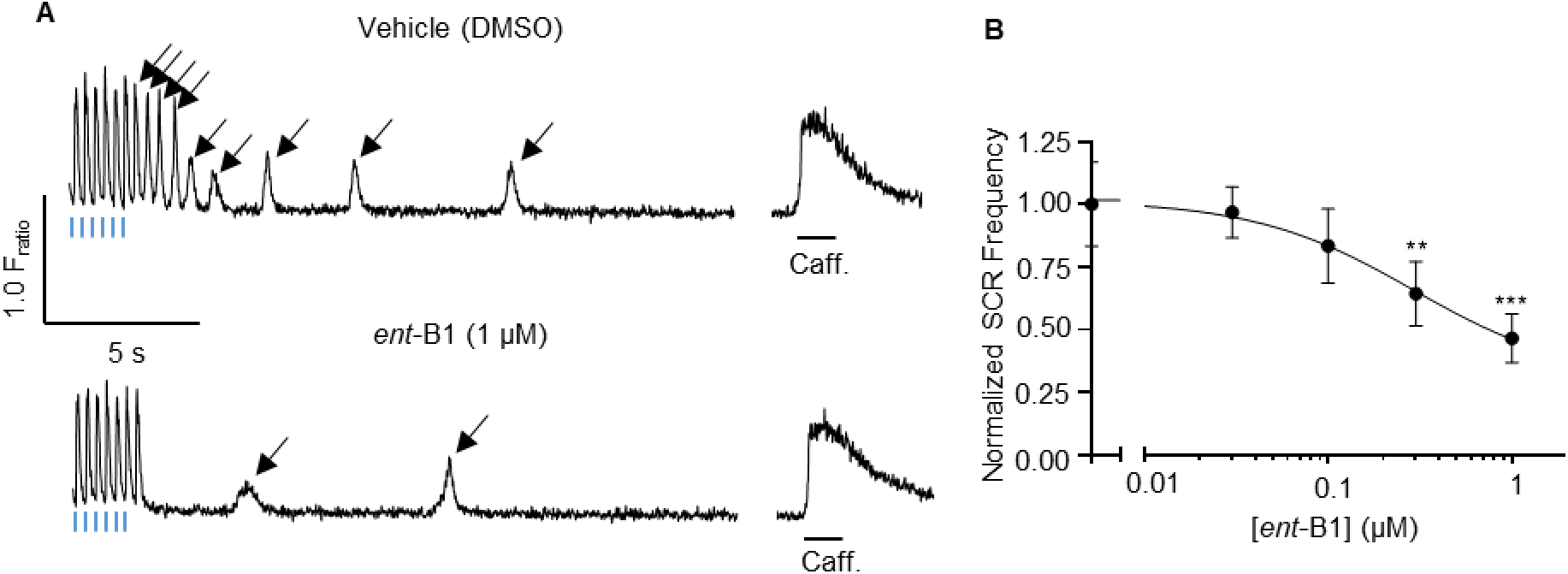
Spontaneous Ca^2+^ release (SCR) in intact mouse *Casq2^-/-^* cardiomyocytes. **A)** Representative recordings from ventricular myocytes isolated from Casq2^-/-^ mice. Cells were field stimulated at 3 Hz for 20 s before recording. Blue marks indicate final stimulations of pacing train, arrows SCR events. Recording duration was 40 s before 10 mM caffeine application to assess total SR Ca^2+^ content. **B)** SCR frequency concentration-response curve for *ent-*B1 following 20 s pacing protocol. *ent-*B1 data normalized to vehicle (DMSO) with 95% CI shown for each concentration. *N* = 26, 30, 30, 30, and 31 cells for 0, 0.03, 0.1, 0.3, and 1 μM *ent*-B1 respectively. Nonlinear regression fit a curve with IC_50_ = 0.23 ± 0.1 μM. ** *P* = 0.0009, *** *P* = 2.4 x 10^-7^ vs DMSO by unpaired, two-tailed *t* test.

### ent-B1 reduces arrhythmia burden in a mouse model of pathologic SR Ca^2+^ release

Once we established that *ent*-B1 could directly bind to RyR2 and permeate the sarcolemma to inhibit pathologic Ca^2+^ release, we wanted to test for therapeutic efficacy in an *in vivo* arrhythmia study. To guide dose selection for efficacy testing, we first established *ent*-B1’s pharmacokinetic properties in mice. Plasma samples were collected and *ent*-B1 concentrations measured after a 3 mg/kg *ent*-B1 i.p. injection. Substantial plasma concentrations were readily achieved, indicating favorable systemic exposure after i.p. administration. The mean peak plasma concentration (C_max_) was 1460 ng/mL at 10 min after injection and exhibited a biphasic decline with a mean elimination half-life (t_1/2_) of 45.4 minutes (**Fig. 4**, **Table 1**). Based on these results, we chose an *in vivo* arrhythmia challenge protocol with the data collection at 15 minutes after *ent*-B1 i.p. administration.

**Figure 4:**
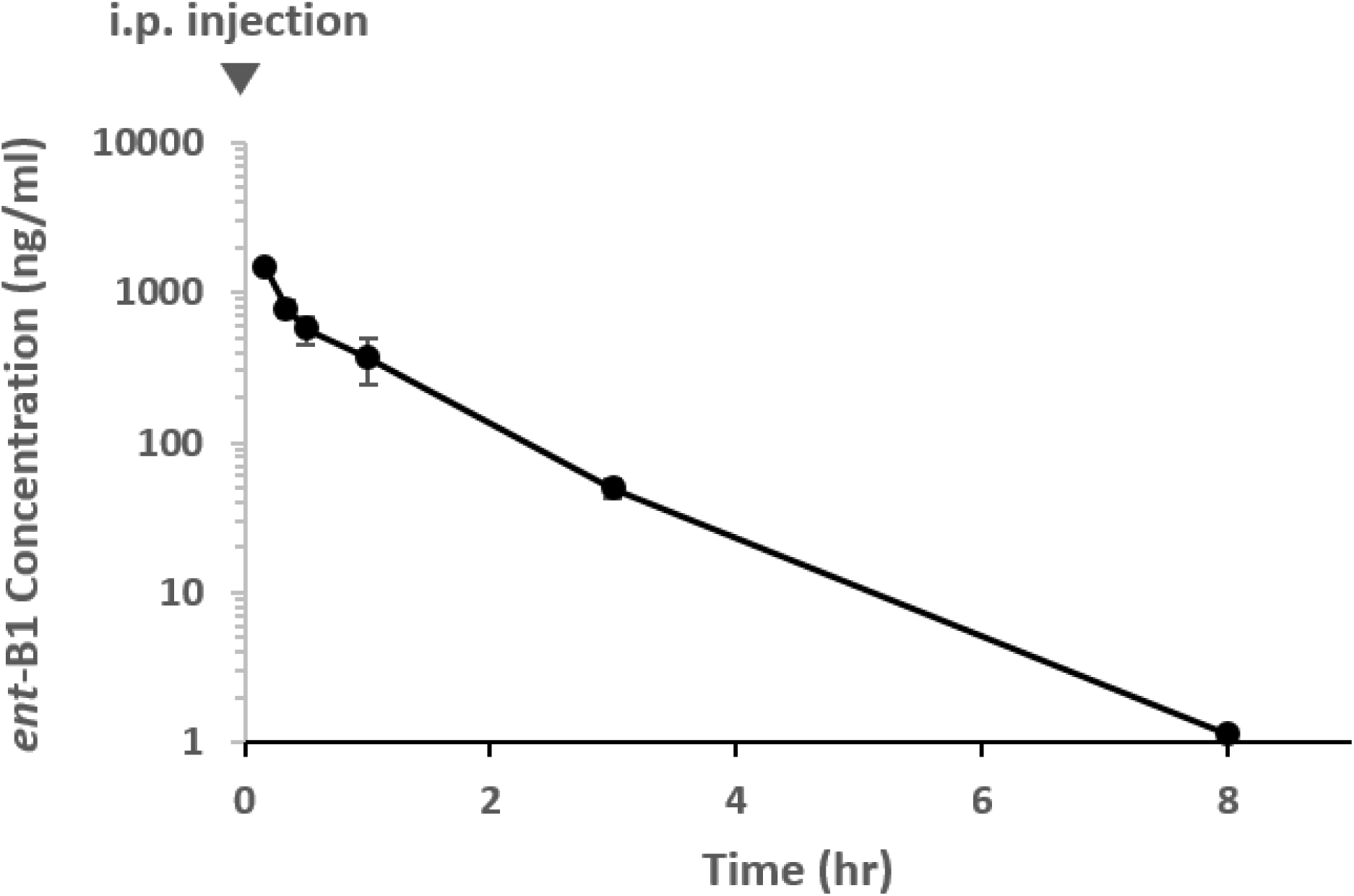
*In vivo ent-*B1 pharmacokinetics in mice. Plasma concentrations of *ent*-B1 after intraperitoneal (i.p.) administration of 3 mg/kg dose. N = 3 mice. Plasma collected serially at 0.167, 0.333. 0.5, 1-, 3-, and 8-hours post-administration (arrowhead, time = 0). Data are mean ± SD with connecting lines.

**Table 1:**
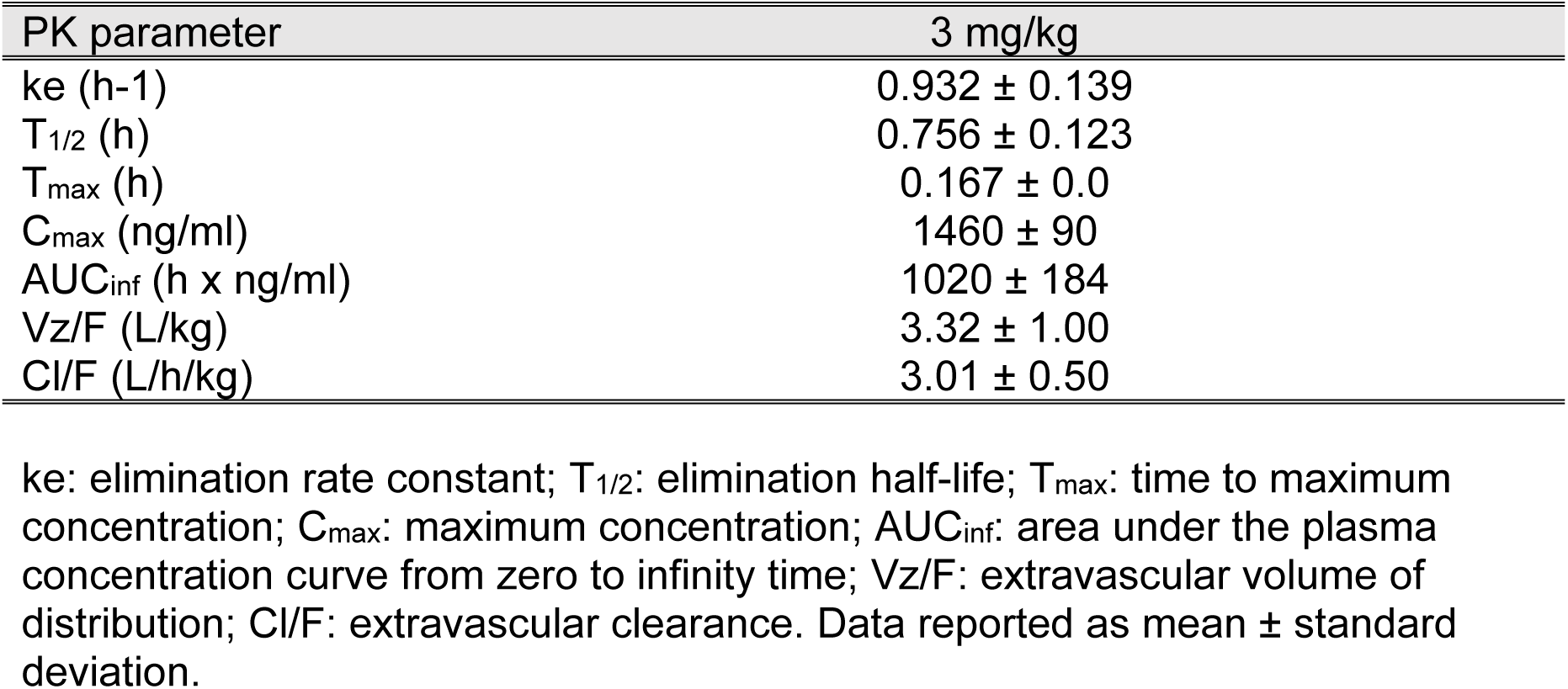
Noncompartmental pharmacokinetic (PK) parameter estimates of ent-B1 following 3 mg/kg intraperitoneal administration in CD1 mice.

*Casq2^-/-^* mice are a validated model for preclinical testing of antiarrhythmic drugs for CPVT (Batiste et al., 2019; Watanabe et al., 2009). Our initial hit compound *ent*-1 had antiarrhythmic efficacy *in vivo* in a single dose study at 3 and 30 mg/kg i.p. in Casq2^-/-^ mice (Blackwell et al., 2023). We tested the same doses of *ent*-B1 in a triple-crossover design with each mouse receiving vehicle (DMSO), 3 mg/kg, and 30 mg/kg with a one-week washout period between experiments. Fifteen minutes after administration, mice were anesthetized with isoflurane and a baseline electrocardiogram (ECG) was established. Ventricular arrhythmias were elicited with a catecholamine challenge using the β-adrenergic agonist isoproterenol (3 mg/kg i.p.), a well-established drug challenge for antiarrhythmic efficacy testing in this model (Watanabe et al., 2009). ECGs were analyzed in blinded fashion and ventricular arrhythmias quantified. **Fig. 5A** gives examples of ventricular arrhythmias induced by the catecholamine challenge. *ent*-B1 caused a dose-dependent reduction in the number of total ectopic beats and incidence of VT (**Fig. 5B, C**). We also scored the mice based on the ventricular arrhythmia severity, i.e., premature ventricular complexes (PVCs) < bigeminy < couplets < ventricular tachycardia (Blackwell et al., 2022). The lower arrhythmia risk scores in mice receiving *ent*-B1 indicate therapeutic efficacy (**Fig. 5D**). No changes in heart rate or QRS width were observed (**Fig. S2**).

**Figure 5:**
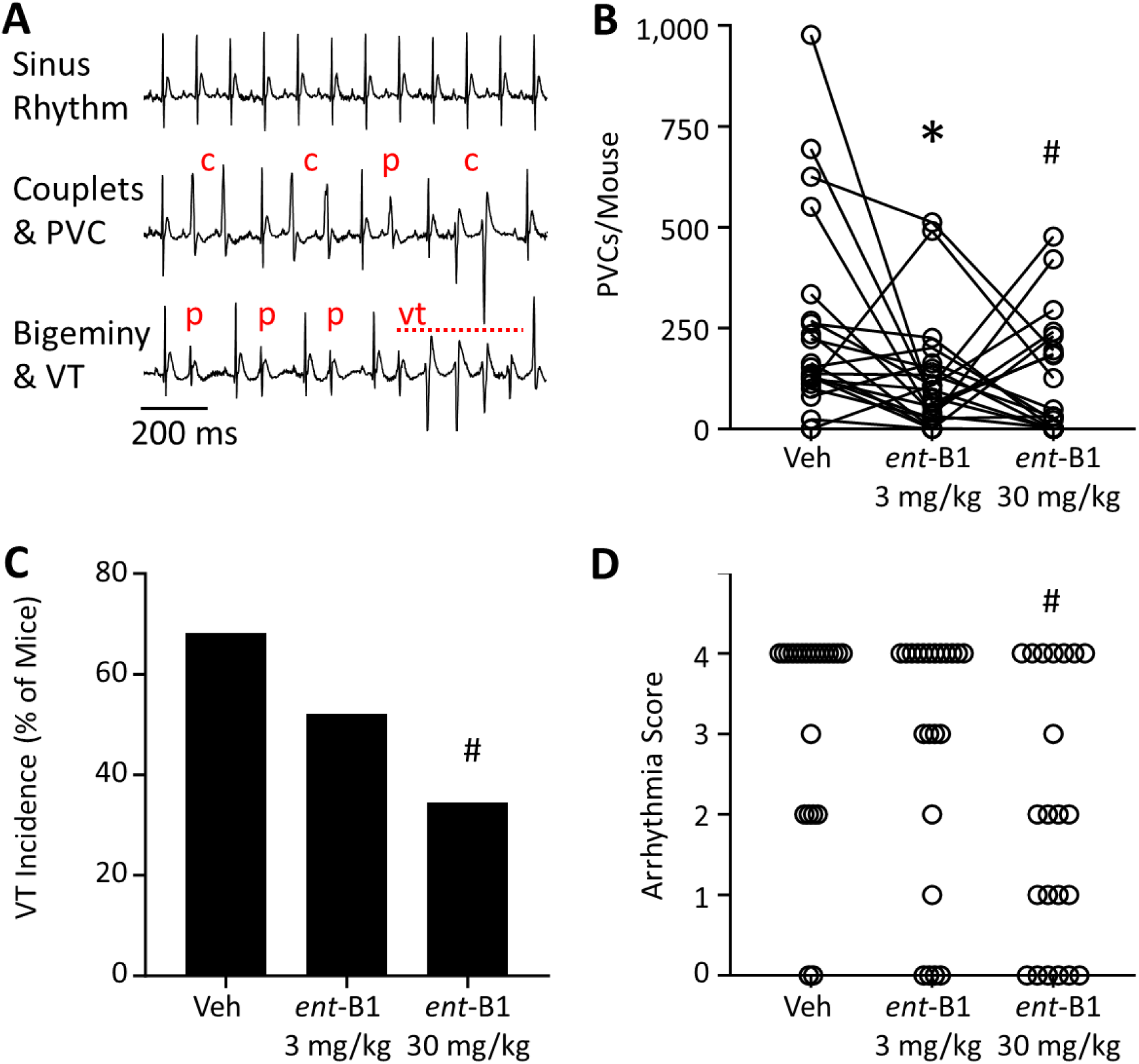
*ent-*B1 antiarrhythmic efficacy in CPVT mice. Triple crossover study design for *in vivo* arrhythmia challenge. N = 22 *Casq2^-/-^* mice were randomly assigned to vehicle (DMSO), 3 mg/kg, or 30 mg/kg *ent-*B1 and crossed over twice with one-week washouts between treatment. Mice underwent catecholamine-induced arrhythmia challenge with isoproterenol 15 min after intraperitoneal drug administration. **A)** Sample ECG traces of normal rhythm (top) and arrhythmias including PVCs and couplets (middle), and bigeminy and ventricular tachycardia (bottom). Ectopic beats are denoted by p (PVC), c (couplet), and vt (ventricular tachycardia). Normal sinus beats are not marked. **B)** Total ectopic beats per mouse including all ventricular arrhythmias (VT). Bonferroni-adjusted p values by pairwise Wilcoxon matched-pairs signed rank test. * P = 0.020 vs Veh; # P = 0.039 vs Veh. **C)** Incidence of ventricular tachycardia. # P = 0.034 vs Vehicle by Fisher’s exact test. **D)** Arrhythmia risk scores were based on an ordinal scale of: 4 = ventricular tachycardia, 3 = couplet, 2 = bigeminy, 1 = isolated PVC, 0 = no PVCs. Bonferroni-adjusted p values by pairwise Wilcoxon matched-pairs signed rank test. # P = 0.0058 vs Vehicle.

## DISCUSSION

Our experiments highlight the potential of leveraging the unique chemical biology of natural product enantiomers to design novel molecular entities with therapeutic efficacy. While the promise of traditional drug candidates rested on maintaining a low molecular weight (<500 Da), limited O and N atoms and H-bond donors as well as hydrophilic properties, the clinical use of natural product compounds and their derivatives is growing despite typically defying one or more of these rules (Caron et al., 2021; Lipinski, 2004; Veber et al., 2002). Within the natural product space, cyclic peptides offer large templates for de novo drug design. For example, we have previously shown that modification of ring size and *N*-methylation of units of *ent*-1 preserve biochemical properties (Smith et al., 2021). Here we show that, in the case of ring size modification, this preserves selectivity and translates to therapeutic efficacy. Further, despite a loss of lipophilicity associated with the smaller compound *ent*-B1 (Smith et al., 2021), there was no apparent impact on membrane permeability. This confirms our hypothesis that the complete structure of the cyclic peptide *ent*-1 is not required for modulation of its target, RyR2.

Our in vitro experiments suggest that *ent*-B1 has a lower potency than *ent*-1. For example, the IC_50_ for *ent*-1 in the ryanodine binding assay is 0.1 µM (Batiste et al., 2019), whereas *ent*-B1 at 1.3 µM (**Fig. 2**) is 10-fold lower. In the intact cardiomyocyte assay, the difference in potency is less, with an IC_50_ of 0.1 µM for *ent*-1 (Batiste et al., 2019), and 0.23 µM for *ent*-B1 (**Fig. 3**), only a 2-fold difference. Although not yet determined, this could indicate that cell membrane diffusion or transport may be faster for *ent*-B1, making it more favorable to engage the intracellular RyR2 target. In fact, although *ent*-1 and *ent*-B1 were not directly compared here, the effect of *ent*-B1 on the clinically relevant sudden cardiac death risk factor VT incidence (∼35%) was comparable to that measured with 30 mg/kg *ent*-1 (∼35%) using the same arrhythmia induction protocol (Blackwell et al., 2023). Hence, *ent*-B1 holds promise as a preclinical candidate given its efficacy in the most physiologically relevant assay, the *in vivo* arrhythmia challenge. Our *in vitro* assays suggest that, like *ent*-1, *ent*-B1 directly binds to RyR2 and is a negative allosteric modulator. *ent*-B1’s combination of submicromolar potency and submaximal efficacy is ideal for our target, as a stronger inhibition of RyR2 could be fatal given its central role in physiologic excitation-contraction coupling. Although peak plasma concentration was comparable to *ent*-1, our pharmacokinetic study revealed a more rapid systemic clearance of *ent*-B1 (t_1/2_ = 45 min, table 1) relative to *ent*-1 (415 min) in mice (Blackwell et al., 2023). Given the short half-life, *ent*-B1 would only be useful in acute clinical scenarios unless the drug formulation is modified to produce a sustained-release profile. Assuming this property is preserved in large animal species and humans, *ent*-B1 could be developed for treatment of ventricular tachycardia storm (“VT storm”), an acute, life-threatening arrhythmia disorder consisting of sequential episodes of sustained VT in a 24-hour window. Taken together, our data with *ent*-B1 further validate RyR2 as an antiarrhythmic drug target and introduce a second preclinical stage compound whose mirror image is a fungal natural product.

## Acknowledgment

This research was supported in part by the National Institutes of Health National Heart, Lung, and Blood Institute [R35 HL144980 (to B.C.K.), R01 HL151223 (to J.N.J., B.C.K., R.L.C.), R01 HL139065 and HL138539 (to R.L.C); the PhRMA Foundation Postdoctoral Fellowship (to D.J.B.); and the American Heart Association Arrhythmia and Sudden Death Strategically Focused Research Network grant [19SFRN34830019 (to B.C.K.)]

## Abbreviations

RyR2: cardiac ryanodine receptor
Casq2: cardiac calsequestrin
CPVT: catecholaminergic ventricular tachycardia
PVC: premature ventricular complex
VT: ventricular tachycardia

## Notes

### Competing Interest Statement

The authors have declared no competing interest.

### Summary of Updates

Corrected spelling, minor text edits to improve flow and updated referencing. No change to results and figures.

