## Supplemental Figures for "*ent*-Verticilide B1 inhibits type 2 ryanodine receptor channels and is antiarrhythmic in Casq2-/- mice"

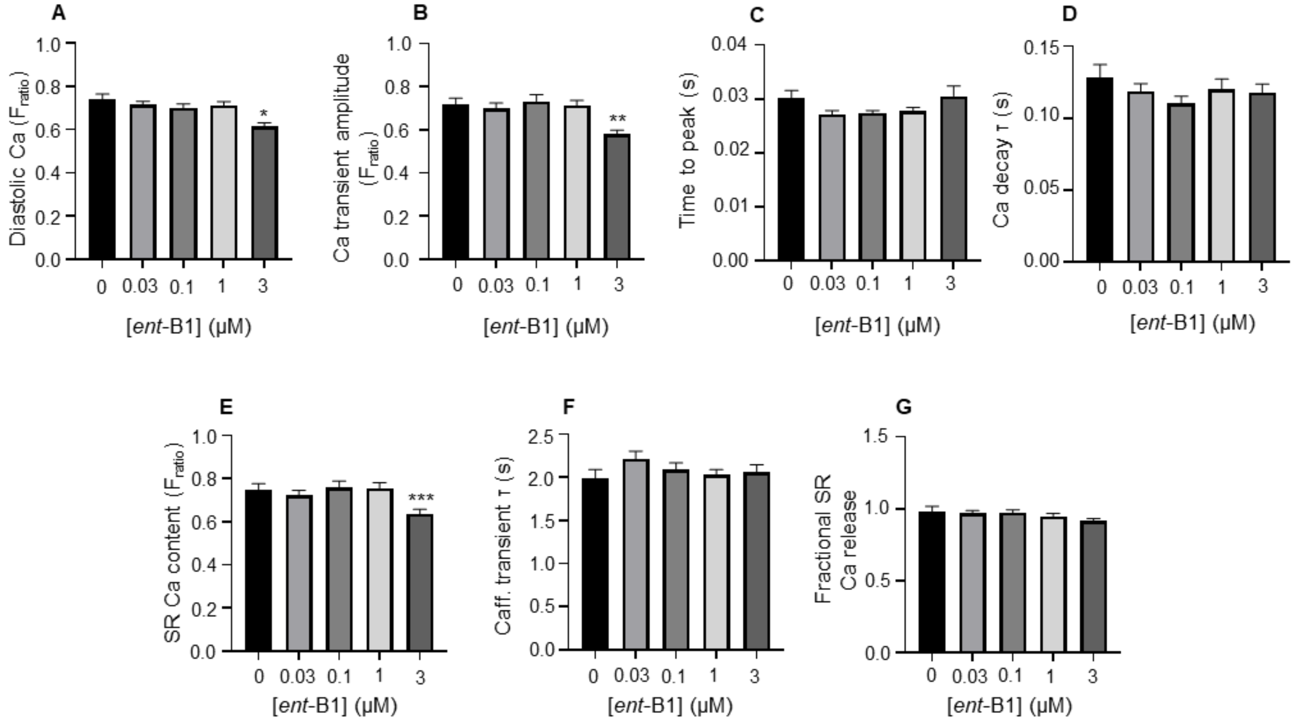


**Supplemental Figure 1: Intracellular $Ca^{2+}$ measurements from intact *Casq2$^{-/-}$* cardiomyocytes. A)** Diastolic $Ca^{2+}$ measured during field stimulation, *P = 3.1 x $10^{-5}$ vs DMSO by t-test. **B)** Paced $Ca^{2+}$ transient amplitudes, **P = 9.6 x $10^{-5}$. **C)** Time to peak and **(D)** $Ca^{2+}$ decay kinetics of paced $Ca^{2+}$ transients**. E**) Caffeine-induced $Ca^{2+}$ transient amplitude, ***P = 0.0024**. F)** Caffeine-induced $Ca^{2+}$ transient decay kinetics. **G)** Fractional $Ca^{2+}$ release = (paced transient amplitude/SR $Ca^{2+}$ content. For panels A-D and G, N = 26, 30, 30, 30, and 31 cells for DMSO, 0.03, 0.1, 1, and 3 μM *ent*-B1, respectively. For panels E and F, N = 14, 27, 29, 29, and 30 cells for DMSO, 0.03, 0.1, 1, and 3 μM *ent*-B1, respectively.

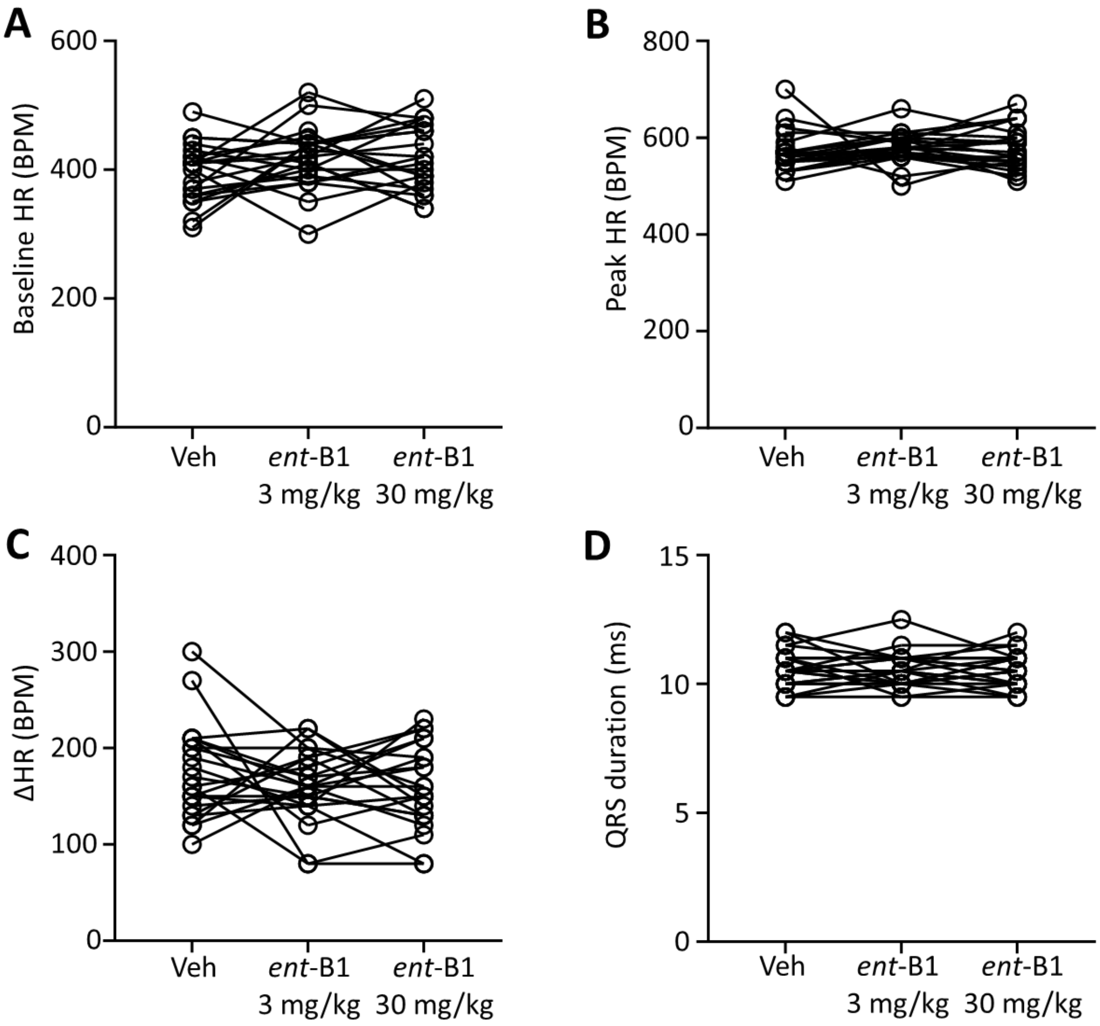


**Supplemental Figure 2**. Electrocardiogram parameters from Casq2-/- mice treated with *ent*-B1. **A)** Baseline heart rate (beats per minute). **B)** Peak heart and **C)** change in heart rate following intraperitoneal administration of 3 mg/kg isoproterenol. **D)** QRS duration at baseline. Mixed effects model pre-test $P > 0.05$ for period, sequence, and treatment in all groups in each panel.
